# Alternative culture systems for bovine oocyte *in vitro* maturation: liquid marbles and differentially shaped 96-well plates

**DOI:** 10.1101/2022.10.24.513502

**Authors:** Andrea Fernández-Montoro, Daniel Angel-Velez, Camilla Benedetti, Nima Azari-Dolatabad, Osvaldo Bogado Pascottini, Krishna Chaitanya Pavani, Ann Van Soom

## Abstract

*In vivo* matured oocytes exhibit higher developmental competence than those matured *in vitro*, but mimicking the *in vivo* environment by *in vitro* conditions has been challenging. Till now, conventional two-dimensional (2D) systems have been used for *in vitro* maturation of bovine cumulus-oocytes-complexes (COCs). However, using such systems may cause cell flattening and does not allow cumulus expansion in all dimensions, which is less physiological. Therefore, implementing a low-cost and highly effective *in vivo*-like microenvironment methodology may help to optimize oocyte *in vitro* maturation. Here, we used two different systems to culture COCs and evaluate their potential influence on embryo development and quality. In the first system, we used treated fumed silica particles to create a 3D microenvironment (liquid marbles; LM) to mature COCs. In the second system, we cultured COCs in 96-well plates with different dimensions (flat, ultra-low attachment round-bottom, and V-shaped 96-well plates). In both systems, the nuclear maturation rate remained similar to the control in 2D, showing that most oocytes reached metaphase II. However, the subsequent blastocyst rate remained lower in the liquid marble system compared to 96-well plates and control 2D systems. Interestingly, a lower total cell number was found in the resulting embryos from both systems (LM and 96-well plates) compared to the control. In conclusion, oocytes matured in liquid marbles or 96-well plates showed no remarkable change in terms of meiotic resumption, embryo development, and quality in both systems. None of the surface geometries influenced embryo development. These findings provide important inferences in many aspects of oocyte and embryo development. Further investigation is needed to determine other aspects like toxicity testing and ultrastructural changes in oocytes.

## Introduction

*In vivo* development of mammalian oocytes depends on continuous contact and communication with cumulus and granulosa cells that are present in the follicle. This interaction results in a dynamic composition of the follicular fluid which is required to fully support the maturation of the oocyte [1,2]. *In vitro* maturation (IVM) systems aims to mimic the *in vivo* environment by using a combination of a base medium supplemented with hormones (FSH, LH), serum or serum replacement, and growth factors [3,4]. However, in the absence of a genital tract, it is often a static system [5,6]. *In vitro* maturation of the oocyte is a crucial step in the acquisition of its developmental competence [7]. Yet, this process is extremely sensitive to environmental factors such as pH [8] or temperature variations [9]. Thus, further optimization of this technique is required to support nuclear and cytoplasmic oocyte maturation and therefore to improve the potential of the oocyte to develop into a blastocyst.

In conventional two-dimensional (2D) systems, cells are grown on flat, firm culture substrates, which are economic and relatively easy to handle. However, in such culture conditions cells can adhere and spread freely in the horizontal plane but they have limited possibility for spreading in the vertical dimension [10]. Hence, the major drawback of 2D-culture systems is that they do not fully imitate the *in vivo* microenvironment where cells are grown in a complex three-dimensional (3D) matrix, which has an impact on cell-cell and cell-extracellular matrix interactions, and consequently on cell responses (differentiation, proliferation, apoptosis, gene, and protein expression) [11–16]. Furthermore, the cell morphology in 2D systems is different from that in the natural structures of tissues, which might affect their functionality, secretion of growth factors, organization of internal structures and cell signaling [17,18].

There is growing evidence suggesting that 3D cell culture models reflect more precisely the actual microenvironment in which cells grow in native tissues [19]. These models allow cell adhesion in all three dimensions whereas in the 2D system it is restricted to the x-y plan. Likewise, 3D systems improve cell communication and soluble factors are more stable in those systems compared to 2D systems [20,21]. In terms of *in vitro* embryo production, studies on 3D systems were designed to support oocyte IVM in several species using alginate microbeads [22], alginate hydrogels [23], glass scaffolds [24], agarose matrix [25], or the hanging drop method [26]. Recently, LM have also become interesting as 3D bioreactors. Liquid marbles are culture medium droplets encapsulated with hydrophobic particles which prevent direct contact between the liquid inside and the surrounding environment, thus, reducing the risk of contamination, while the hydrophobic shell of the LM remains permeable for gases [27]. These properties make LM a promising alternative as 3D microbioreactors for cell culture. This system has been used for culturing microorganisms [28], embryoid bodies [29,30] or olfactory ensheathing cell spheroids [31]. Liquid marbles have been also applied in ovine [32] and porcine [23] oocyte IVM but it has not been validated yet in the bovine model.

Cellular responses are influenced by topographical surface features. For instance, human epithelial cells presented differences in orientation, migration, and morphology when culturing them on pillar or pit surfaces [33]. Similarly, concave and convex surfaces have been found to influence stem cells' differentiation into osteoblasts [34], and V-shape surface has been related to changes in cell shape and mRNA expression in fibroblasts [35,36] and osteoblast-like cells [37]. However, all these studies used complex systems to recreate the different surface geometries. Thus, a simpler alternative could be to use 96-well plates with different shapes, which are available on the market and are easy to use and standardize. Three different 96-well plates have been tested in several human cell lines (retinal epithelial, alveolar epithelial and dermal fibroblastic) [38], nevertheless, to our knowledge, no studies have been performed with these plates to evaluate the potential influence of different surface topographies during oocyte IVM on embryo development.

In the present work, we evaluated the potential effects of using a 3D culture system during oocyte IVM with two different methods: (1) a matrix system, using liquid marble microbioreactors and (2) a non-matrix system, using differently shaped culture substrates (flat, round, and v-shaped 96-well plates). We found both matrix and non-matrix 3D culture systems had a similar effect as 2D culture systems in terms of oocyte nuclear maturation, while embryo development was similar after oocyte maturation in the 96-well plates but lower in LM.

## Materials and methods

### Experimental design

#### Experiment 1: Evaluation of liquid marble as 3-D model for *in vitro* maturation

Liquid marbles were tested as microbioreactors for *in vitro* maturation of bovine oocytes. To do so, a total of 941 cumulus-oocyte complexes (COCs) in seven replicates were used in three different maturation systems: Liquid Marbles (LM) (n= 301 COCs) as 3D model, 2D droplets (n = 309 COCs) as flat culture with similar oocyte/medium ratio to LM (5 COCs/30 μL), and Control group (n = 331 COCs) as our standard condition (60 COCs /500 μL). After maturation, COCs from all groups were randomly distributed for *in vitro* fertilization (IVF) and *in vitro* culture (IVC; LM = 241, 2D droplets = 256, and Control = 262 COCs) or nuclear maturation assessment (LM = 60, 2D droplets = 53, and Control = 69 COCs).

#### Experiment 2: Evaluation of different surface geometries for *in vitro* maturation

A comparative study was carried out to evaluate the effect of three different bottom-shaped multi-well plates during oocyte IVM on embryo development. Firstly, we conducted a pilot study including 3 replicates (n = 1,414 COCs) to select the ideal work volume of maturation medium per well and to analyze the effect of paraffin oil overlay in oocytes matured in multi-well plates. Cumulus-oocyte complexes were matured in v-shaped 96-well plates under the following conditions: five COCs in 30 μL maturation medium with 30 μL oil overlay (Oil V-shaped-30 (OV-30); n = 160 COCs) or without oil (V-shaped-30 (V-30); n = 120 COCs), ten COCs in 60 μL maturation medium with 30 μL oil overlay (OV-60; n = 163 COCs) or without oil (V-60; n = 155 COCs), and twenty COCs in 120 μL maturation medium with 30 μL oil overlay (OV-120; n = 143 COCs) or without oil (V-120; n = 148 COCs). A control group (n = 252 COCs) was also included. After maturation, IVF and IVC were performed routinely. Subsequently, based on the results of the pilot study, three different surface geometries (flat (F), ultra-low attachment round-bottom (R), and V-shaped (V) 96-well plates) were compared during maturation in six replicates (n = 1,992 COCs) using the conditions described as for the OV-60, resulting in 4 groups: F-60 (n = 427 COCs), R-60 (n = 549 COCs), V-60 (n = 491 COCs) and control group (n = 525 COCs). After maturation, COCs from all groups were randomly assigned to IVF and IVC (F-60 = 374, R-60 = 504, V-60 = 440 and control group = 487 COCs) or nuclear maturation assessment (F-60 = 53, R-60 = 45, V-60 = 51 and control group = 38 COCs). The experimental design is depicted in Fig 1.

**Figure 1.**
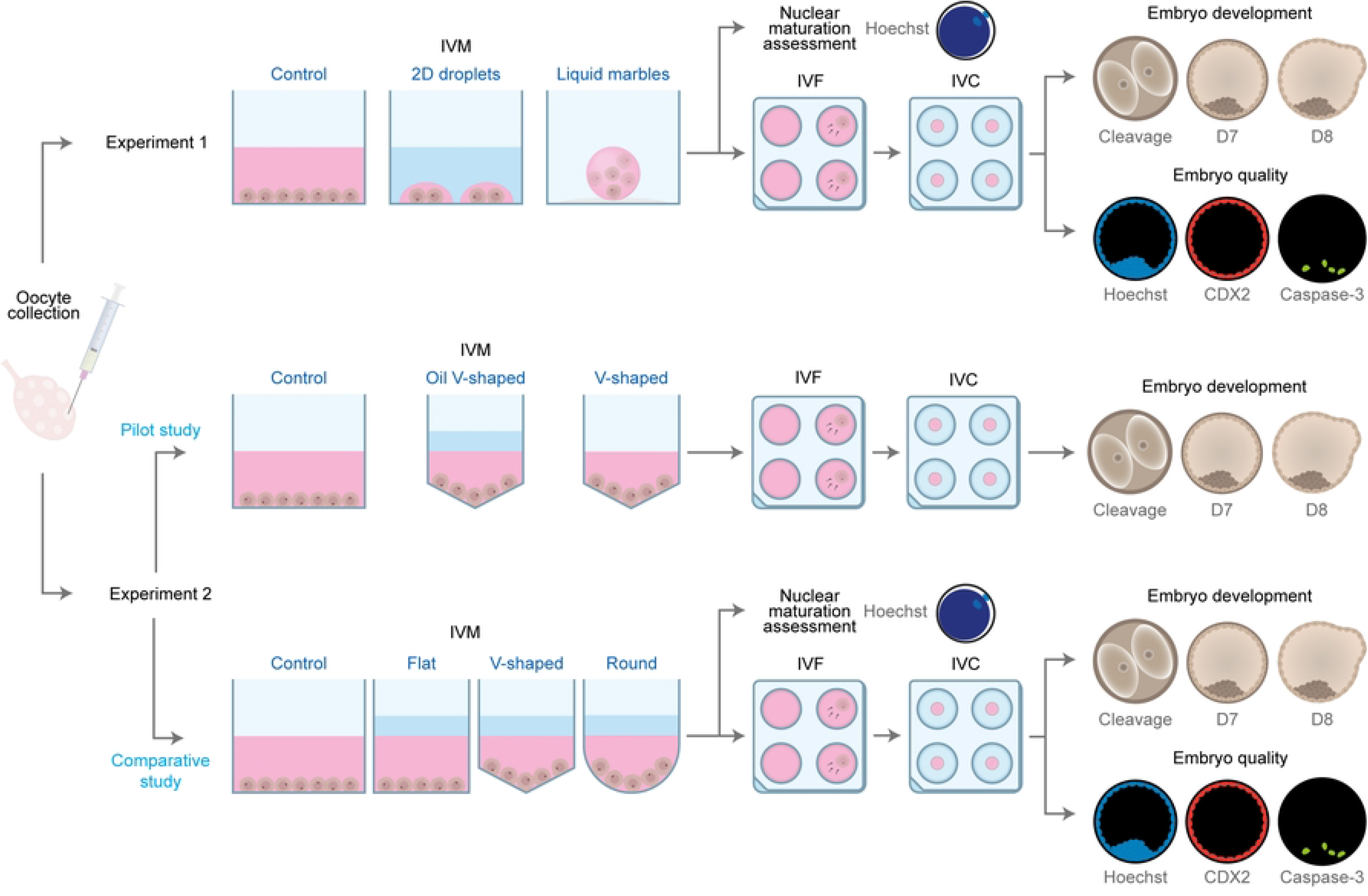
Schematic representation of the experimental design.

### Media and reagents

Tissue culture media (TCM)-199 and antibiotics (gentamycin and kanamycin) were obtained from Life Technologies Europe (Ghent, Belgium). Phosphate-Buffered Saline (PBS) was purchased from Gibco™ 20012019, Thermo Fisher Scientific (Waltham, MA, USA). All other products not indicated otherwise were provided by Sigma-Aldrich (Diegem, Belgium). Before use, every media were filtered (0.22 μM; GE Healthcare-Whatman, Diegem, Belgium).

### Source of oocytes and *in vitro* maturation

Bovine ovaries were obtained from a local slaughterhouse, transported to the laboratory, and prepared for further processing within 2 h after collection. The ovaries were disinfected with 96% ethanol and cleaned for three times in physiological saline (37 °C) containing 50 mg/mL of kanamycin. Cumulus-oocyte complexes were recovered with an 18-gauge needle and a 10 mL syringe from 4-8-mm-diameter follicles. Oocytes surrounded by three or more layers of compact cumulus cells and a uniformly granulated cytoplasm were selected, washed in warm HEPES - Tyrode’s Albumin Lactate Pyruvate media (HEPES-TALP), and randomly assigned to different IVM systems. Four IVM systems, as described below, were evaluated according to the experimental group (see Experimental design). All treatment groups were cultured for 22 h in 5% CO_2_ in the air at 38.5 °C.

#### Control

Sixty COCs were cultured in 500 μL maturation medium (TCM-199 Earle’s salts supplemented with 20 ng/mL epidermal growth factor and 50 μg/mL gentamicin) in flat-bottom 4-well dishes (Thermo Fisher^®^, Merelbeke, Belgium) without oil covering.

#### Encapsulation in liquid marbles

A single droplet of 30 μL maturation medium containing five COCs was carefully placed on top of a layer of approximately 1 cm treated fumed silica powder (Cabot Corp, Cab-O-Sil, TS-530), which was equally distributed in a 6 cm Petri dish (Fig 2A). The Petri dish was mildly shaken in circular motions to ensure that the surface of the droplet was completely and uniformly coated with the hydrophobic particles. To manipulate the LM, the edge of a 1000 µl micropipette tip was cut to make its diameter to a small extent of the LM diameter in order to ensure a proper grip, but big enough to avoid collapse. Before transferring the marbles, the modified tip was coated with some powder to prevent its adhesion to the tip. Then, the LM was picked up slowly (Fig 2B) and placed on a well of a 24-well plate (Thermo Scientific) whose surface was previously covered with a small quantity of silica powder (Fig 2C). To avoid evaporation, the central space of the 24-well plate was filled with 5 mL sterile HEPES-TALP medium. After IVM, the LM (Fig 2D-2E) was placed in maturation medium to disrupt the silica powder's hydrophobicity, causing the marble's dissolution (Fig 2F). The released COCs were washed three times in maturation medium to remove silica particles before proceeding to the next step.

**Figure 2.**
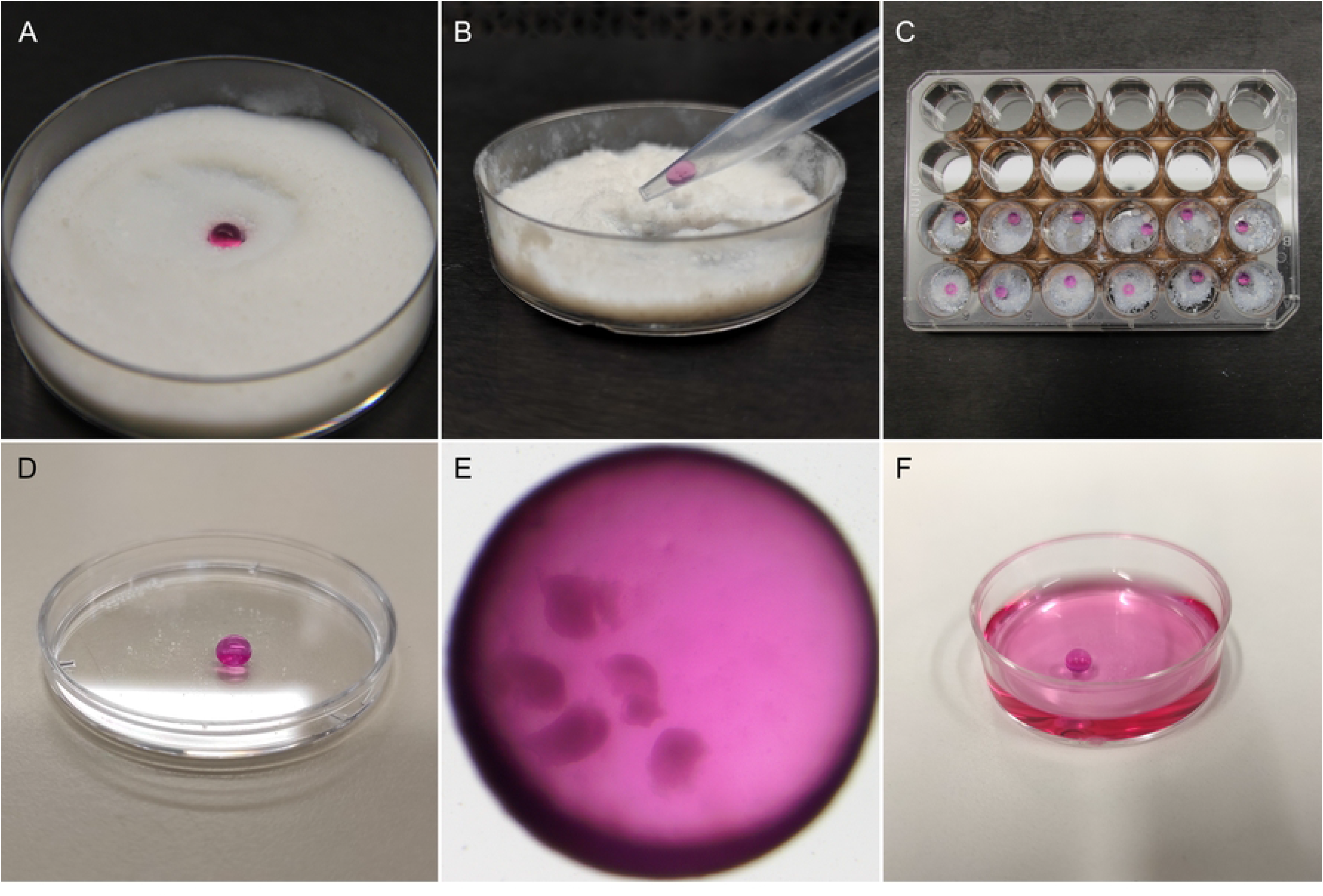
Oocyte encapsulation in liquid marbles. (A) One droplet of maturation medium containing the oocytes was placed in treated fumed silica powder on a petri dish. The petri dish was gently shaken to form the liquid marble. (B) A modified 1000 µl micropipette tip was used to manipulate the liquid marbles. (C) Liquid marbles were placed individually in the wells of a 24-well plate containing a small amount of silica powder. The central space of the plate was filled with 5 mL HEPES-TALP to prevent evaporation. (D) Resulting liquid marble drop. (E) Five COCs encapsulated in a liquid marble drop before IVM, observed under a stereomicroscope. (F) After maturation, liquid marbles were dissolved in maturation medium.

#### 2D droplets

Droplets of 30 μL maturation medium were prepared in a Petri dish (60 × 15 mm; Thermo Fisher Scientific, Waltham, MA USA) and covered with 7.5 mL paraffin oil (SAGE, CooperSurgical, Trumbull, CT, USA). Five COCs were matured in each droplet of maturation medium.

#### Shaped culture in 96-well plates

Firstly, oocytes were matured in V-shaped 96-well plates under the following conditions: five COCs in 30 μL of maturation medium with or without paraffin oil overlay of 30 μL, ten COCs in 60 μL of maturation medium with or without a paraffin oil overlay of 30 μL, and twenty COCs in 120 μL of maturation medium with or without a paraffin oil overlay of 30 μL. Secondly, ten COCs were matured in 60 μL of maturation medium with paraffin oil overlay in flat, ultra-low attachment round-bottom, and V-shaped 96-well plates (all from Corning^®^, Houten, Netherlands).

### *In vitro* fertilization and embryo culture

Standard *in vitro* methods were used to generate bovine embryos, as previously described by Wydooghe et al. [39]. Briefly, using a discontinuous 45/90% Percoll^®^ gradient (GE Healthcare Biosciences, Uppsala, Sweden), sperm capacitation of frozen-thawed straws from a known fertile bull was performed. Consequently, the sperm pellet was washed in IVF–TALP medium and a final concentration of 1 × 10^6^ spermatozoa/mL was adjusted using IVF–TALP medium enriched with BSA (Sigma A8806; 6 mg/mL) and heparin (25 mg/mL).

After 22 hours of IVM, oocytes from each treatment group in experiments 1 and 2 were pooled to reach groups of 60 COCs. Then, oocytes were washed in IVF-TALP and subsequently co-incubated in 500 μL IVF-TALP with Percoll-purified spermatozoa for 21 h at 38.5 °C in 5% CO_2_ in humidified air. After fertilization, zona attached sperm and cumulus cells were removed by vortexing for 3 minutes in 2.5 mL Hepes-TALP. The presumed zygotes were randomly selected and cultured in groups of 25 in 50 μL droplets of synthetic oviductal fluid (SOF), 0.4% (w/v) BSA (Sigma A9647), and ITS (5 μg/mL insulin, 5 μg/mL transferrin, and 5 ng/mL selenium). Each droplet was covered with 900 μL paraffin oil and incubated at 38.5 °C for 8 days in 5% CO_2_, 5% O_2_, and 90% N_2_.

Cleavage was evaluated 45 h post insemination and blastocyst yield was recorded on day 7 and day 8 post insemination. Both rates were calculated as a percentage over the presumed zygotes.

### Evaluation of oocyte nuclear stage (maturation assessment)

After maturation, oocytes were denuded by vortexing for 8 min in 2.5 mL Hepes-TALP and fixed with 4% paraformaldehyde (w/v). Then, oocytes were transferred to 0.1% (w/v) polyvinylpyrrolidone (PVP) in PBS containing 10 μg/ml of Hoechst 33342 (Life Technologies, Ghent, Belgium) for 10 min. Nuclear morphology was evaluated using a fluorescence microscope (BRESSER Science ADL 601 F LED). The proportion of oocytes in each meiotic stage – germinal vesicle (GV), germinal vesicle breakdown (GVBD), metaphase I (MI), metaphase II (MII), or degenerated – was recorded.

### Embryo quality assessment

Embryo quality was determined by differential apoptotic staining for CDX2, a transcription factor only expressed by trophectoderm cells, and caspase-3, a cysteine-aspartic acid protease involved in the signaling pathways of cell apoptosis. The protocol was performed accordingly to Wydooghe et al. [40]. Briefly, day 8 blastocysts were fixed in 4% paraformaldehyde (w/v) at room temperature for at least 20 min and then stored in PBS supplemented with 0.5% BSA at 4 °C until the staining was performed. Firstly, blastocysts were incubated with ready-to-use anti-CDX2 primary antibodies (Biogenex, San Ramon, USA). Embryos were next incubated with rabbit active caspase-3 primary antibody (0.768 ng/mL, Cell Signaling Technology, Leiden, The Netherlands), followed by incubation in goat anti-mouse Texas Red secondary antibody (20 μg/mL in blocking solution, Molecular Probes, Merelbeke, Belgium) and then in goat antirabbit FITC secondary antibody (10 μg/ml in blocking solution, Molecular Probes). Finally, the embryos were transferred to nuclear stain, Hoechst 33342 (50 μg/mL in PBS/BSA). A negative control was also included in which embryos were not incubated with CDX2 and active caspase-3 antibodies. Samples were examined by a single observer using fluorescence microscopy (Leica DM 5500 B) with a triple bandpass filter. With this staining protocol, the number of trophectoderm (TE) cells, inner cell mass number (ICM), total cell number (TCN = TE + ICM), ICM/TCN ratio, the total number of apoptotic cells (AC) and the ratio of apoptotic cells (ACR; AC/TCN) were estimated.

### Statistical Analyses

The statistical analyses were performed using R-core (version 4.2.1; R Core Team, Vienna, Austria). The oocyte/zygote/embryo was considered as the unit of interest. Generalized mixed-effects models were used to test the effect of IVM conditions on oocyte nuclear maturation, cleavage, and embryo development rates. The effect of IVM conditions on blastocyst differential staining parameters was fitted in mixed linear regression models. For all the models, the replicate was set as random. Results are expressed as least square means and standard errors. The differences between treatment groups were assessed using Tukey’s post hoc test. The significance and tendency levels were set at p < 0.05 and p < 0.1, respectively.

## Results

### Experiment 1: Evaluation of liquid marbles as a 3D model for *in vitro* maturation

#### Effect of liquid marbles on oocyte nuclear maturation

Nuclear maturation assessment by Hoechst staining demonstrated that oocytes in the three groups resumed meiosis (i.e., no germinal vesicles were found). Most oocytes in all treatment groups reached the metaphase II stage with no significant differences among groups (p > 0.05; Table 1). Likewise, the proportion of oocytes that reached germinal vesicle breakdown, metaphase I, or degenerated was similar among groups (p > 0.05).

**Table 1.**
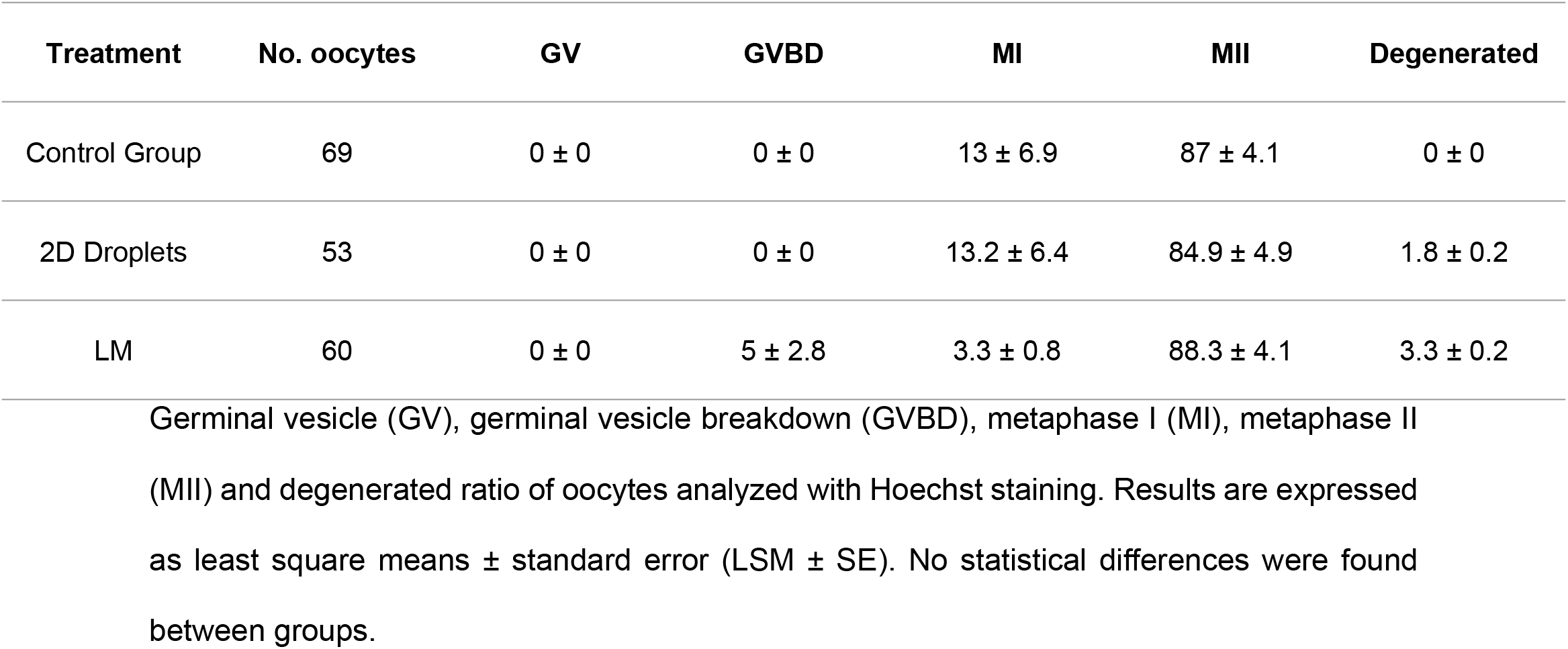
Nuclear maturation assessment of oocytes matured in: (A) control group, (B) 2D droplets, and (C) liquid marbles (LM).

| Treatment | No. oocytes | GV | GVBD | MI | MII | Degenerated |
| --- | --- | --- | --- | --- | --- | --- |
| Control Group | 69 | 0 ± 0 | 0 ± 0 | 13 ± 6.9 | 87 ± 4.1 | 0 ± 0 |
| 2D Droplets | 53 | 0 ± 0 | 0 ± 0 | 13.2 ± 6.4 | 84.9 ± 4.9 | 1.8 ± 0.2 |
| LM | 60 | 0 ± 0 | 5 ± 2.8 | 3.3 ± 0.8 | 88.3 ± 4.1 | 3.3 ± 0.2 |

Germinal vesicle (GV), germinal vesicle breakdown (GVBD), metaphase I (MI), metaphase II (MII) and degenerated ratio of oocytes analyzed with Hoechst staining. Results are expressed as least square means ± standard error (LSM ± SE). No statistical differences were found between groups.

#### Effect of liquid marbles on embryo development and embryo quality

Firstly, we determine the effect of oocyte maturation conditions on cleavage and blastocyst rates. Although there were no significant differences in the cleavage rates between groups (LM: 79.9 ± 3.1%; 2D droplets: 85.7 ± 2.9%; control group: 86.0 ± 3.1%; p > 0.05; Fig 3A), oocytes matured in LM showed lower day 7 (17.6 ± 3.4%) and day 8 (26.1 ± 3.7%) blastocyst rates compared to 2D droplets (26.4 ± 4.2%, p = 0.048, and 38.8 ± 4.2%, p = 0.008, respectively) and control (29.8 ± 4.9%, p = 0.01, and 40.1 ± 4.6%, p = 0.007, respectively).

**Figure 3.**
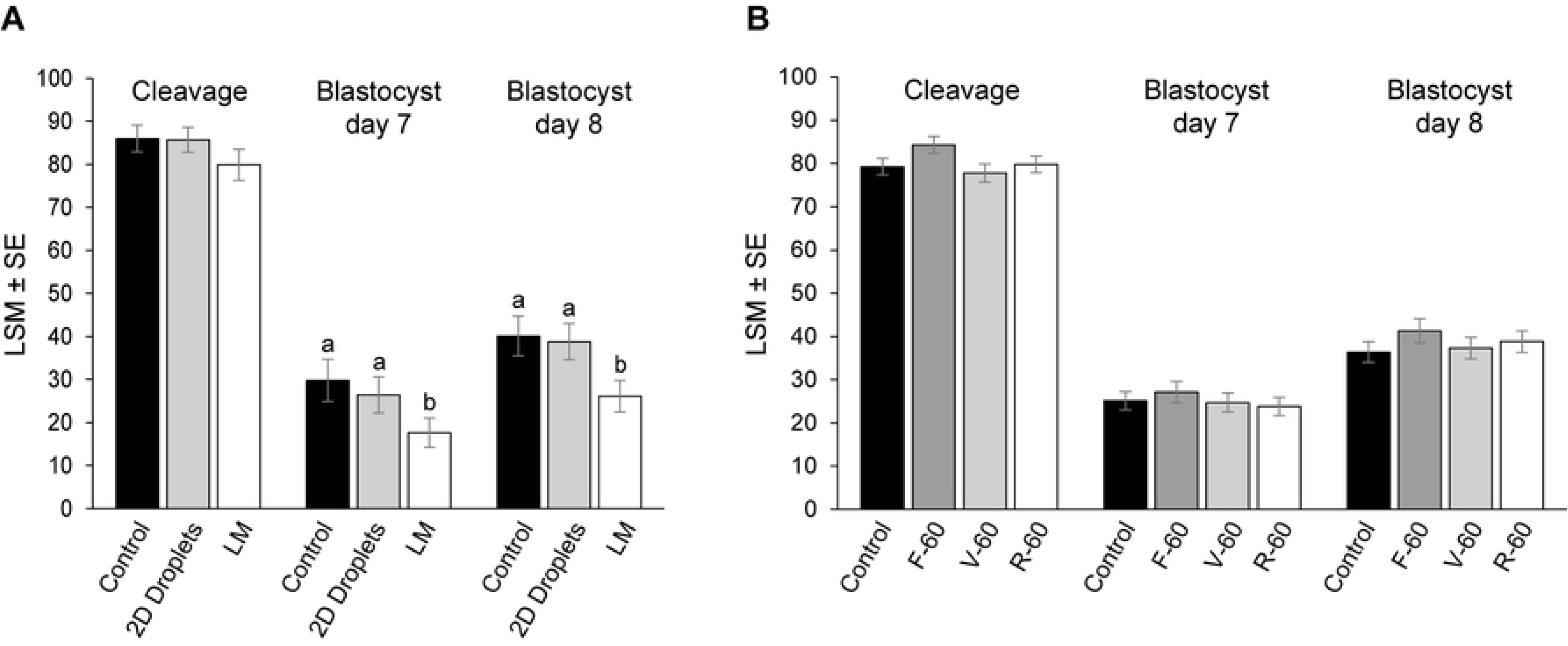
Cleavage, day 7, and day 8 blastocyst rates are expressed as a percentage of presumed zygotes. (A) Experiment 1. Oocytes were *in vitro* matured in liquid marbles (LM), 2D droplets, and a control group. (B) Experiment 2. Oocytes were *in vitro* matured in flat, v-shaped, and ultra-low attachment round-bottom 96-well plates, and a control group. Different superscripts (a and b) represent statistical differences (p < 0.05) among groups. Results are expressed as least square means ± standard error (LSM ± SE).

Further on, we determined the effect of maturation conditions on embryo quality by differential apoptotic staining of blastocysts. Differences among IVM culture systems in blastocyst quality parameters are shown in Table 2. Maturation in LM produced blastocysts with lower TCN and TE than in control (p < 0.01). Maturation in 2D droplets reduced the TCN, ICM, and TE (p < 0.01) and increased the AC/TCN ratio compared to control (p = 0.03).

**Table 2.** Effect of the oocyte *in vitro* maturation in LM on embryo quality.

| Treatment | No. blastocyst | Cell numbers |  |  |  | ICM/TCN | AC/TCN |
| --- | --- | --- | --- | --- | --- | --- | --- |
|  |  | TCN | ICM | TE | AC | ratio | ratio |
| Control Group | 52 | 117.8 ± 8.6<sup>a</sup> | 36.1 ± 2.2<sup>a</sup> | 81.6 ± 4.0<sup>a</sup> | 1.9 ± 3.7 | 30.9 ± 1.3 | 2.0 ± 0.5<sup>a</sup> |
| 2D Droplets | 51 | 82.3 ± 5.1<sup>b</sup> | 26.2 ± 2.2<sup>b</sup> | 56.3 ± 4.0<sup>b</sup> | 2.8 ± 3.7 | 32.6 ± 1.4 | 3.9 ± 0.5<sup>b</sup> |
| LM | 53 | 91.3 ± 5.1<sup>b</sup> | 31.6 ± 2.2<sup>ab</sup> | 59.8 ± 4.0<sup>b</sup> | 3.0 ± 3.6 | 33.7 ± 1.3 | 3.5 ± 0.5<sup>ab</sup> |

Total cell number (TCN), trophectoderm cells (TE), inner cell mass (ICM), apoptotic cells (AC), ICM/TCN ratio, and AC/TCN ratio of day 8 blastocyst differentially stained. Different superscripts per column (a and b) represent statistical differences ( $p < 0.05$ ) among groups. Results are expressed as least square means ± standard error (LSM ± SE).

### Experiment 2: Evaluation of different surface geometries for *in vitro* maturation

Initially, a pilot study was performed to establish the optimum volume of maturation medium and the effect of paraffin oil overlay during oocyte IVM in 96-well plates on embryo development. The V-30 group was excluded from the treatment groups due to excessive maturation medium evaporation after IVM. Although there were no significant differences in the cleavage, blastocyst day 7 and blastocyst day 8 rates for all the treatment groups compared to the control (p > 0.05), OV-60 had a numerically higher blastocyst rate at day 8. Therefore, we selected IVM culture conditions as described for this group for follow-up experiments. For detailed information on the pilot study results, see supporting information (S1 Table).

### Effect of surface geometry on oocyte nuclear maturation, embryo development, and embryo quality

Hoechst staining was used to analyze the meiotic progression of the oocytes after IVM in flat, v-shaped, and ultra-low attachment round-bottom 96-well plates. No differences were found in the proportion of mature, immature, or degenerated oocytes in the three tested geometries and the control group (p > 0.05; Table 3).

**Table 3.** Nuclear maturation assessment of oocytes matured in: (A) control group, (B) F-60, (C) V-60, and (D) R-60.

| Treatment | No. oocytes | GV | GVBD | MI | MII | Degenerated |
| --- | --- | --- | --- | --- | --- | --- |
| Control Group | 38 | $2.6 \pm 0$ | $7.9 \pm 4.4$ | $2.6 \pm 0.1$ | $86.8 \pm 5.5$ | $0 \pm 0$ |
| F60 | 53 | $3.8 \pm 0$ | $11.3 \pm 4.4$ | $0 \pm 0$ | $84.9 \pm 4.9$ | $0 \pm 0$ |
| V60 | 51 | $0 \pm 0$ | $5.9 \pm 3.3$ | $3.9 \pm 0.1$ | $90.2 \pm 4.2$ | $0 \pm 0$ |
| R60 | 45 | $4.4 \pm 0$ | $4.4 \pm 3.1$ | $0 \pm 0$ | $91.1 \pm 4.2$ | $0 \pm 0$ |

Germinal vesicle (GV), germinal vesicle breakdown (GVBD), metaphase I (MI), metaphase II (MII) and degenerated ratio of oocytes analyzed with Hoechst staining. Results are expressed as least square means ± standard error (LSM ± SE).

Cleavage, day 7 and day 8 blastocysts rates were similar in the three 96-well plates and the control group (p > 0.05; Fig 3B). Differences among treatments in blastocyst quality parameters determined by differential apoptotic staining are shown in Table 4. *In vitro* maturation in flat, ultra-low attachment round-bottom and v-shaped 96-well plates resulted in blastocysts with lower TCN and TE than control (p < 0.05). Blastocysts in the R-60 group also presented lower ICM compared to control (p = 0.01). However, there were no differences in the ICM/TCN or AC/TCN ratios among treatments and control (p > 0.05).

**Table 4.**
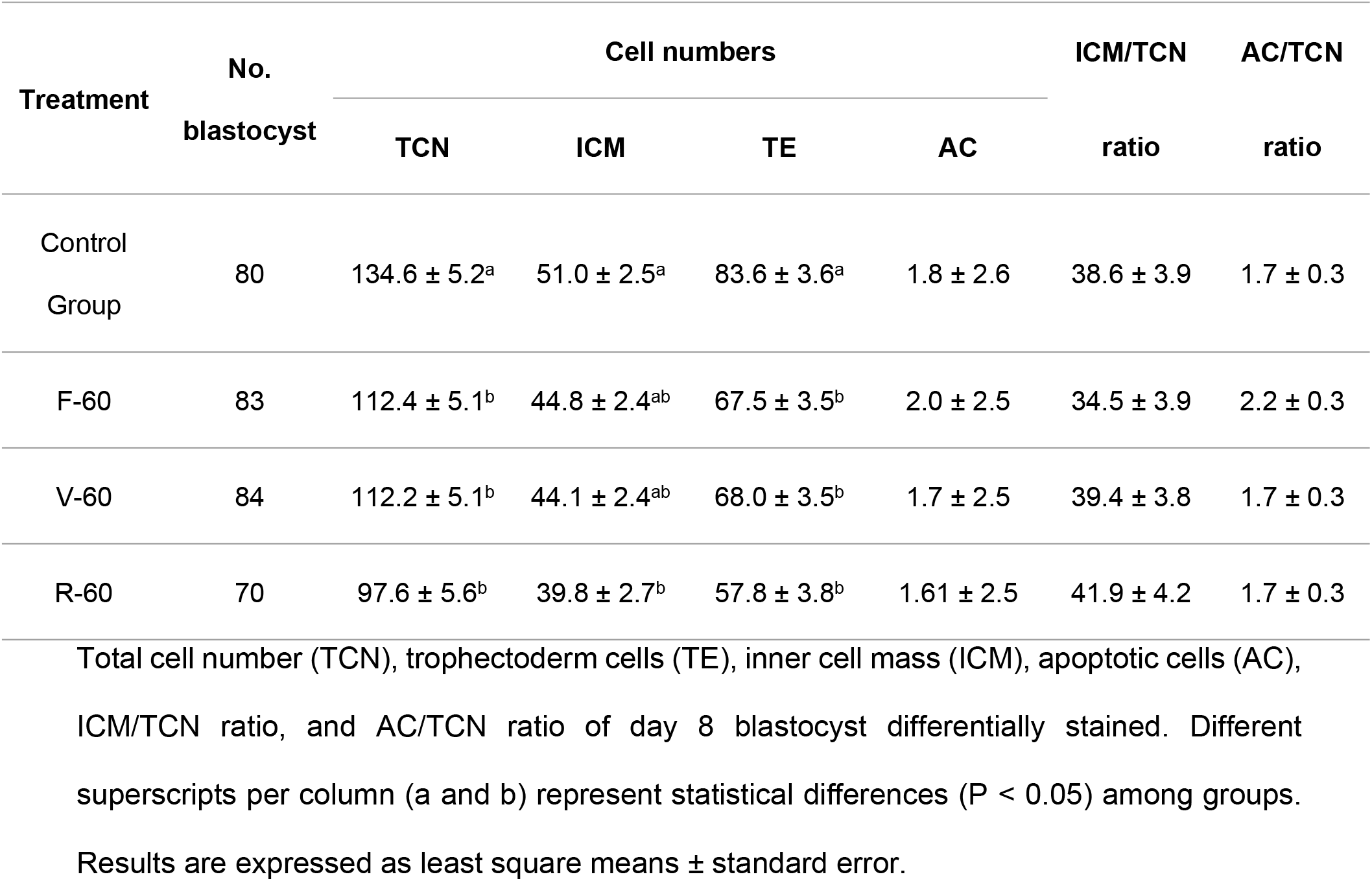
Effect of the oocyte *in vitro* maturation in three different surface geometries on embryo quality.

## Discussion

Currently, most standard oocyte IVM is performed in two-dimensional culture systems, which are both economical and practical. However, these systems are only a poor representation of the physiological environment within the follicle. Therefore, novel alternatives to mature oocytes in a three-dimensional milieu might enhance the interaction between the oocyte, the cumulus cells, and different factors within the culture medium, and consequently improve the developmental competence of the oocyte. In this study, we used liquid marbles as microreactors to perform serum-free IVM for the first time in the bovine model. Moreover, we also tested a simple and practical non-matrix system using differently shaped 96-well plates. We found that both matrix and non-matrix 3D culture systems had a similar effect as 2D culture systems in terms of oocyte nuclear maturation, while embryo development was similar after oocyte maturation in the 96-well plates but lower in LM.

We proved that meiotic resumption is not affected in bovine oocytes by the use of LM, since the proportion of oocytes that reached metaphase II was similar in both 2D- and 3D-systems, which concord with previous studies in the cat [41] and sheep [32,41]. In our study, although oocytes matured in LM were able to reach the blastocyst stage, the blastocyst yield was reduced compared to oocytes matured in 2D. However, embryos derived from oocytes matured in LM exhibited a lower total cell number count than those natured in a 2D system. It has been demonstrated that material toxicity can decrease the total cell number and affect embryo rates [42]. However, the toxicity of treated fumed silica particles has not been tested in oocytes. Therefore, we hypothesize that treated fumed silica particles could exhibit some toxicity in bovine oocytes since our findings differ from the results obtained by Bebbere et al., who matured ovine oocytes in LM formed with the same particles, showing better blastocyst rate compared to those matured in 2D conditions [41]. Additionally, in a previous study of the same group, better blastocyst yield was reached after the maturation of ovine oocytes in polytetrafluoroethylene marbles compared to the group in 2D [32]. Therefore, apart from the possible toxicity, the presence of serum in their maturation medium, the oocyte/medium ratio, or species-specific differences might explain discrepancy of the results. Moreover, additional manipulation during the LM preparation compared to the standard system may also affect the developmental capacity of the gametes.

Besides the LM system, in the current study, we evaluated the effect of different geometry surfaces on oocyte developmental competence. Although we tested a range of medium volumes and evidenced the importance of paraffin oil overlay when low volumes are used, we did not find differences in oocyte nuclear maturation nor embryo development, or quality between the three surface geometries tested. Although blastocyst derived from oocytes matured in 96-well plates presented a lower total cell number count than oocytes matured in standard conditions, culture conditions did not adversely affect the developmental capacity of the oocytes. These results are in agreement with Shafaie et al., the only previous study that compared flat, round-bottom, and V-shaped 96-well plates in cell culture. In that study, different human cell lines, namely A549 (alveolar epithelial), ARPE-19 (retinal epithelial) and Malme-3M (dermal fibroblast), cells attached and spread differently in each plate, but the phenotype and functionality of the cells were not affected by the surface topography [38].

As cell culture is tending to move towards more physiological systems such as 3D-methods, it is important to study the impact of those systems on fundamental cellular processes. In *in vitro* embryo production, oocyte maturation and embryo development and quality are the main outcome parameters to evaluate culture systems. In our study, we showed that oocyte meiotic resumption was not affected by any of the culture systems. However, only nuclear maturation was assessed and further studies to evaluate differences in cytoplasmic maturation may be performed. Interestingly, our experiments with different surface topographies exhibited a similar blastocyst rate in comparison with traditional culture but LM reduced it. Yet, alternative culture systems entail higher costs and complexity than conventional IVM (Table 5). Liquid marbles demonstrated the highest level of complexity and risk of loss of oocytes during the elaboration of the marble, with the longest handling time, which might be an additional explanation for the lower blastocyst yield. On the other hand, 96-well dishes made the handling of COCs during preparation and recovery more difficult due to the smaller diameter of the wells.

**Table 5.** Characteristics of *in vitro* maturation systems.

| Maturation technique | Description | Hands-on-time (min) |  | Ease-of-use | Cost (€) |
| --- | --- | --- | --- | --- | --- |
|  |  | Preparation | Recovery |  |  |
| Four well dish without oil (control) | Sixty COCs in 500 μL of maturation medium in flat-bottom 4-well dishes without oil covering | 1 | 0.5 | + | 1.3 |
| Droplets on petri dish under oil (2-D Droplets) | Five COCs in droplets of 30 μL of maturation medium in a Petri dish (60 x 15 mm) and covered with 7.5 mL paraffin oil | 4 | 3 | ++ | 3.2 |
| Liquid marbles | Five COCs in 30 μL of maturation medium are placed on top of a layer of treated fumed silica powder to form a LM, which is transferred to a well of a 24-well plate with a cut 1000 μL pipette tip | 15 | 7 | ++++ | 3.3 |
| 96-well plates with different dimensions | Five COCs in 60 μL of maturation medium covered by 30 μL of paraffin oil overlay in flat, ultra-low attachment round-bottom, and V-shaped 96-well plates | 4 | 5 | +++ | Flat: 3.8<br>V-shaped: 3.9<br>Round-bottom: 22.3 |

Comparison of *in vitro* maturation systems in terms of working principle, hands-on-time (i.e. based on the manipulation time in minutes required to handle 60 COCs from the dish in which they are selected and washed in HEPES-TALP to the final *in vitro* maturation (IVM) system (preparation) and from the IVM system to the fertilization dish (recovery)), ease-of-use and approximate cost for one maturation process (i.e. based on cost of the dish, oil if necessary and silica powder for liquid marbles preparation). + = low; ++ = moderate; +++ = high; ++++ = very high

## Conclusion

For the first time, two different 3D culture systems were tested in bovine IVM. We showed that there are limited differences in development and quality in embryos resulted from oocytes cultured in the 2D control system compared with LM and 96-well plate systems. Importantly, no adverse results were observed in terms of embryo development when cultured in 96-well plates and, although there was a slight decrease in embryo yield in LM, these results contribute towards the development of a more accurate, simple high throughput, *in vitro* maturation systems that represent the *in vivo* dynamic environment. Maybe further investigation is needed like in toxicity testing and ultrastructural changes of the oocyte.

## Acknowledgments

The authors thank Petra Van Damme for her technical assistance. This project has received funding from the European Union’s Horizon 2020 research and innovation programme under the Marie Skłodowska-Curie grant agreement No 860960.

